# An Examination of the Use of Large Language Models to Aid Analysis of Textual Data

**DOI:** 10.1101/2023.07.17.549361

**Authors:** Robert H. Tai, Lillian R. Bentley, Xin Xia, Jason M. Sitt, Sarah C. Fankhauser, Ana M. Chicas-Mosier, Barnas G. Monteith

**Affiliations:** University of Virginia; Oxford College of Emory University; University of Kansas; THInc AI Group

**Keywords:** Qualitative methodology, Large Language Models (LLMs), Deductive qualitative coding, Reliability

## Abstract

The increasing use of machine learning and Large Language Models (LLMs) opens up opportunities to use these artificially intelligent algorithms in novel ways. This article proposes a methodology using LLMs to support traditional deductive coding in qualitative research. We began our analysis with three different sample texts taken from existing interviews. Next, we created a codebook and inputted the sample text and codebook into an LLM. We asked the LLM to determine if the codes were present in a sample text provided and requested evidence to support the coding. The sample texts were inputted 160 times to record changes between iterations of the LLM response. Each iteration was analogous to a new coder deductively analyzing the text with the codebook information. In our results, we present the outputs for these recursive analyses, along with a comparison of the LLM coding to evaluations made by human coders using traditional coding methods. We argue that LLM analysis can aid qualitative researchers by deductively coding transcripts, providing a systematic and reliable platform for code identification, and offering a means of avoiding analysis misalignment. Implications of using LLM in research praxis are discussed, along with current limitations.

## Introduction

Text is an essential form of qualitative data, and its analysis enables researchers to understand complex social phenomena (Dey, 2003). Traditionally, qualitative analysis of text involves meticulous, time-consuming, and labor-intensive manual coding. This subjective process depends on several factors of a given researcher’s background and experience, i.e., positionality, which may introduce bias in the analysis and interpretation of qualitative data. To mitigate bias, reliability measures often involve two or more researchers analyzing, i.e., “coding,” the data. The themes, concepts, and phenomena that are identified are referred to as “codes.” Similarities and differences in the coding can be identified, and inter-rater reliability can be calculated (e.g. Cohen, 1960; Fleiss, 1971). Recent advancements in natural language processing and machine learning have opened up new possibilities for assisting qualitative researchers in their analytical endeavors (Manning, 2022). In this paper, we outline a systematic methodology for using Large Language Models (LLM) as a qualitative research tool that can aid in the deductive coding of interview data.

Qualitative codes serve as interpretive labels or categories describing various themes, concepts, or phenomena in the textual data. These codes can be applied in an inductive or deductive manner, depending on the researchers’ positionality and role in their work (Elo & Kyngäs, 2008). Inductive coding requires the researcher to read the textual data and identify codes within the text. This allows the researchers to build patterns from the data, eventually organizing it into a comprehensive list of themes. Researchers apply a series of codes to the text for deductive analysis based on a theoretical framework. When performing deductive coding, researchers often follow a detailed “codebook” to ensure the coding matches predetermined characteristics or theoretical constructs that are being applied (Schwandt, 1997). This paper will focus on the use of an LLM in deductive coding.

Coding can be subjective, mainly when only one researcher performs a textual analysis. Unintended bias may creep into an analysis in several ways. How a researcher approaches that data or how they define particular terms present in the text may have an influence on the findings of a researcher. This subjectivity can lead to variations in how researchers interpret and assign codes to the data (Marques & McCall, 2005).

Some researchers create memos, writing and reflecting on their thoughts and ideas to address potential researcher bias. If multiple researchers are coding the same data, they can apply an inter-rater statistic to quantify the degree of similarity between coders (Marques & McCall, 2005) When coders disagree, negotiations occur between them to refine the coding and reach an agreement. Codebooks are commonly re-defined and re-applied to the text. This iterative process helps focus the coding and increases reliability, but is also highly labor intensive and depends on researcher subjectivity.

This research article aims to examine the systematic use of an LLM as a qualitative analysis tool that can supplement the deductive coding process. An LLM is a type of artificial intelligence (AI) algorithm. It is a trained deep-learning model that can understand and generate text, answer questions, and complete other language-related tasks with high accuracy (Kasneci et al., 2023). Some examples include ChatGPT^®^, Open AI^®^, and Bing Chat^®^. By leveraging contextual understanding and LLMs, researchers may verify and enhance alignment in qualitative textual analysis, potentially improving the overall quality of the output.

## Literature Review

To lay the groundwork for using an LLM as a deductive qualitative analysis tool, we review how LLM works to create sizeable textual data networks and discuss some of the problems that researchers should consider when using LLMs. We then consider *deductive coding in the qualitative analysis of narrative text, outline some of the current reliability measures of traditional coding use* and make a case for the addition of LLM as a new research instrument. Our focus will shift to recent empirical work, which uses ChatGPT^®^ 3.5 as an exemplar. ChatGPT^®^ 3.5 is one of several available LLMs and was chosen for its ubiquity, wide recognition, and accessibility. However, the reader should be aware that the analytical method we outline in this paper is applicable across all current forms of LLMs and is not limited to ChatGPT^®^. Our analysis will anchor the discussion in the current academic landscape. Opportunities for methodological expansion using LLMs as a reliability measure in qualitative research will also be discussed.

### LLM and Data Processing

LLMs comprise large computing systems called Artificial Neural Networks (ANN). Textual data that are inputted into the ANN are transformed into numerical data to be later processed by the central processing unit of the computer. As more data are input into the LLM, the ANNs become more powerful and form interconnected layers of information (Valdenegro, 2023). LLMs like ChatGPT^®^ 3.5 have been trained to model human preference, learning from human feedback (Wang et al., 2021). ChatGPT® 3.5 can conduct fluent conversations with people and solve mathematical problems (Frieder et al., 2023), and has been found to possess a range of emergent abilities (Wei et al., 2022). These capabilities have attracted widespread public attention and provide an opportunity to explore how ChatGPT^®^ 3.5 can be used in data analysis procedures.

One issue with LLMs and data processing is hallucinations. Hallucinations are a phenomenon where the LLM generates nonsensical information or an illogical output (Athaluri et al., 2023). Generative AI follows a complex algorithm that responds to human input rather than deciding what is true. Likewise, these hallucinations can arise from adversarial examples, which include diverse input data that can perplex LLM systems. This conflicting information can lead to misclassification and misinterpretation, producing inaccurate and “hallucinatory” outputs (Athaluri et al.). To combat the production of hallucinations, it is essential to create contextually-relevant prompts and to include human reviewers to assess outputs generated by LLM systems. In this article, we do not argue that LLMs should replace human coding but instead highlight the possibility of using LLMs as an additional tool for validation in qualitative research.

### Deductive Coding and Reliability

One popular approach to qualitative analysis in social science is using deductive codes. In deductive coding, researchers develop a qualitative manual that contains a list of predetermined definitions that researchers apply to their data. Frequently, these codes are derived from theoretical frameworks and are used to test the presence or absence of specific characteristics or themes within the text. The codebook typically provides a list of codes, a label for each code, brief definitions, and an example of a quote that illustrates the use of each code (Creswell & Creswell, 2017).

To ensure the reliability of deductive coding, researchers can calculate different reliability statistics. For example, Cohen’s (1960) kappa statistic can be used to measure the level of agreement between two raters. Cohen’s kappa quantifies the extent to which raters agree on the relative ratings or coding and serves as a measure of accuracy (Linacre, 1989). Another example of an inter-rater statistic is Fleiss’ kappa, which is used when there are more than two coders (Fleiss, 1971). Since Fleiss’ kappa is a derivative of Cohen’s kappa, it also uses the hypothetical probability of agreement by chance. Each of these measurements attempts to ensure the reliability of data analysis by comparing sets of codes to each other.

Although Cohen’s and Fleiss’ kappa provide quantifiable statistics for qualitative research, there are limitations to applying these measures. For example, Block and Kraemer (1989) argued that because Cohen’s kappa allows marginal distributions to differ, it measures the association between two sets of ratings instead of the agreement between two raters. Nicolas et al. (2011) argued that Fleiss’ kappa does not apply weighting during analysis, which results in variation between coders for the same dataset, even if there is no variation. Using LLMs as another comparison tool could help qualitative researchers expand upon the different reliability measures.

### LLMs and Empirical Research

The use of LLMs is being adopted in a variety of fields. For example, ChatGPT^®^ has been used in translation and text generation across linguistic research fields (Jiao et al., 2023; Peng et al., 2023; Ubani & Nielsen, 2023). It has also been used in the medical field to write discharge summaries (Patel & Lam, 2023), to help radiologists in their decision-making processes (Rao et al., 2023), and to help diagnose prostate cancer (Van Booven et al., 2021). This broad range of applications highlights the potential to use ChatGPT^®^ in novel ways.

Focusing on academic writing, Bhardwaz and Kumar (2023) evaluated using two LLMs, Google Bard® (Mountain View, CA) and ChatGPT®, in writing literature reviews. The authors aimed to determine how well the LLMs could summarize the text without plagiarism. The authors asked Google Bard^®^ and ChatGPT^®^ to paraphrase the abstracts of ten articles. They demonstrated that texts written by the study authors had low plagiarism rates, while texts written by Google Bard^®^ and ChatGPT^®^ had comparatively much higher plagiarism counts. Similarly, Rahman et al. (2023) used ChatGPT^®^ 3.0 to write an academic paper. They discovered that ChatGPT^®^ 3.0 could generate problem statements and an outline, but artificial intelligence could not draft a validated literature review or quantify numerical data. The references for the literature review section of the academic paper were cited correctly using APA formats, but the citations were not connected to authentic empirical articles. The authors concluded that LLMs could be used to analyze qualitative data but did not expound upon this potential application.

Narrowing the focus on qualitative research, Xiao et al. (2023) used ChatGPT^®^ 3.0 to investigate the potential use of LLMs for qualitative analysis. According to the authors, ChatGPT^®^ 3.0 can be used to code data deductively, and this coding is comparable to traditional coding when the LLM is given a clear codebook. They also analyzed different prompt designs that were input into ChatGPT^®^ 3.0. They concluded that codebook-type prompts that provide structure and contextual information are more reliable than example-type prompts, those that provide information without additional context. However, presenting the codebook and at least one example prompt produced the most accurate results compared to traditional coders. This work lays the foundation for exploring different applications of deductive coding with LLM.

There is a paucity of validated research examining the use of LLMs for qualitative analysis, specifically with multiple iterations of text being put into ChatGPT® 3.5 to test the consistency of results. A benefit of using LLMs in qualitative analysis is that it provides almost unlimited inter-rater measures. Each time a researcher logs into an LLM network, the prompt entered into the model becomes a new input text, and this action is analogous to a new rater coding the data. The LLM incorporates some randomness in the way it processes and weighs information. This randomness is measured and defaults to 1 (relatively random) in ChatGPT®. This random integration results in responses that may not agree with prior outputs (Gilardi et al., 2023). Researchers can then evaluate whether the imputed codebooks and examples sufficiently meet the assumptions of kappa statistics or if additional codebooks and examples are needed. This improves the efficiency of the researcher’s time, focusing on accurately depicting the coded data and reducing the personnel needed for reliable evaluations.

### Purpose

To date, no developed systematic practices exist for using LLMs for qualitative data analysis. As LLMs evolve and presumably become more reliable, it is crucial to explore analytical practices and protocols as qualitative research applications based on LLMs arise. This work aims to outline a new method for using LLMs that may contribute to the validity and reliability of qualitative analysis techniques of textual data. In this paper, we address the following research questions:

### Research Questions

1. How does the output of a large language model respond to the recursive entry of a qualitative data analysis prompt across a range of iterations?
2. How do the responses of a large language model compare to the traditional qualitative analysis on the same three textual data sources?

## Methodology

The focus of this study was to develop a procedure for using an LLM to analyze interview-based textual data and to test the reliability of the procedure. This methodology section will outline the participants that were part of the study, the sample used to run the iterations of codebook analysis in ChatGPT^®^ 3.5, the data collection techniques, and data analysis.

### Data Source

The data used in this study were drawn from archival interviews from *Project Crossover: A Study of the Transition from Student to Scientist* (NSF REC 0440002), a sequential, mixed-methods study exploring the transition of Ph.D. students to independent researchers. This project examined the experiences of individuals who had or were at the time, engaged in the process of becoming a scientific researcher in chemistry, physics, or chemical engineering (Dabney & Tai, 2014). As part of the project, 125 semi-structured interviews were collected from chemistry, physics, and chemical engineering graduate students, postdocs, scientists, research engineers, and individuals who had left the field of scientific research. Each interview was recorded and transcribed. They ranged in length from 30 minutes to 2.5 hours.

### Sample

This study used three excerpts from two Project Crossover interviews. These interviews captured these scientists’ beliefs of what characteristics constitute a successful scientist. Interviews focused on graduate school, pre-graduate educational history, and research and employment experiences. Project Crossover interviews were additionally screened for relevance based on the identification of five characteristics, which are referred to as codes:

1. autonomy or self-motivation to pursue research experiences,
2. persistence despite adversity or difficulty,
3. perception of researcher identity,
4. the desire to create novel knowledge, and
5. interest in engaging in research in a STEM-related discipline.

### Data Collection

#### ChatGPT^®^ 3.5

We used ChatGPT^®^ 3.5 by OpenAI^®^ because of its general ubiquity. The reader should be aware that while the analysis in this paper uses ChatGPT^®^ 3.5, the authors are not advocating for using this platform over any other LLM. The techniques we explore in this study are generally applicable across several LLM platforms such as BingChat^®^ by Microsoft^®^, Bard AI^®^ by Google^®^, Claude 2^®^ by Antropic^®^, LLaMa 2^®^ by Meta^®^, and ErnieBOT^®^ by Baidu^®^, among others. ChatGPT^®^ 3.5 is an autoregressive language model with more than 175 billion parameters, 10x more than any previous non-sparse language model (Brown et al., 2020). As ChatGPT^®^ evolves online, variations are released, and the LLM becomes faster and more refined. At the time of writing, the accessible version of ChatGPT^®^ was built on OpenAI^®^’s Generative Pre-Trained Transformer (GPT) 3.5. GPT 4 is available via a paid subscription. In this study, all references to ChatGPT^®^ are limited to version 3.5 unless otherwise noted. For readers seeking a complete comparison of different LLMs and their capacities, a table addressing these comparisons is provided in Nepo et al., 2017 and a comparative review is provided in Ray, 2023.

#### Codes and Prompts

For this study, we examined five different codes: *Autonomy*, *Persistence*, “*Perception of Own Identity and Self*,” *Novelty*, and *STEM Interests*. The research team inductively created these codes through the analysis of science fair participant interviews that were collected and analyzed for a separate research project. The definitions of the codes are shown below. To combat hallucinations, we created contextually relevant prompts with clear definitions and kept the structure of prompts identical for each iteration. We designed the LLM prompt for a “binary” query (i.e., a yes-or-no answer). If the LLM produced a vague response to the query, we took a conservative approach and coded it as “no.” For the first query to the LLM, we entered the prompt and interview sample, Text 1, below.

We defined the characteristics below:

1. Autonomy: is the ability to be self-driven in STEM research
2. Persistence: is the continuance in a course of action in spite of difficulty
3. “Perception of own identity or self”: is the internal recognition as an individual who can talk about research with an expert in the discipline or confidence
4. Novelty: is the desire to create new knowledge or knowledge new to them
5. STEM interests: is having an interest in a STEM field.

Can you find the five characteristics in the transcript below, yes or no? If yes, give us the quote.

*[Text 1]*

*It really was, I guess, working with the postdoc. I, of course, tried to be as independent as I could be and I did try a number of things, but then I would brainstorm with the postdoc and we met often with the advisor. I’m actually a very organized person and so if I can get a list of six things to try, I can work through them. So it was with the help of others*.

For the second and third queries, we replaced Text 1, shown in italics above, with Text 2 and Text 3, respectively.

[Text 2]

In several ways. One, again, at the University, you’re a scientist educator. And so that aspect of it, being successful is having my graduate students enjoy, have a passion for science, and find rewards in it, helping them to learn to think and analyze and become scientists of their own. That’s a success, teaching the next generation. And being a successful scientist—wow. I mean I think it would be wonderful to be able to really make a major advance in terms of medicine or some really breakthrough area, but for me it’s probably just discovering little bits about Mother Nature and helping it fit into the big picture. What I mentioned about the biochemistry earlier, in many ways I would love to be able to move a little bit that way where I think the ability to effect the well-being of people is stronger….Well, I certainly intelligence and creativity are necessary. More and more, a strong work ethic helps. I think an ability to communicate with others, both students and other scientists because, often, again, and what we were talking in the graduate school, that often we learn by interacting with others. So, yeah, that’s certainly main, I mean, certainly a passion for the work. Science is, especially in academics now, one of the most highly paid professions, and so it really does take someone who is committed, who just really enjoys science.

[Text 3]

Whereas the veteran researcher is happy to be confused, because if you’re not confused you’re not going to learn anything that’s worthwhile, because it must be already understood. And if you’re confused there’s a chance that when you finally figure out what’s going on there’s a chance you will have learned something that’s of interest to other researchers. So that’s the general thrust of what I have in mind.…It’s a question of how thoughtful and original, and I say spunky or enterprising and all the students are. I cultivate their maturity, that sort of thing. If they, for example, show initiative in going to the literature, or going to talk to somebody else at another university, or somewhere, or finding something on the web, and then getting in touch with someone else all on their own initiative, it shows that they’re really totally thinking of this as their thing, and carrying the ball and so forth….I think the kind of science I strive to do is that sort of spirit. It’s a question of can you do something as an architect that opens people’s eyes to new possibilities and gets them excited. I like to tell my students that the real value of an experiment is spiritual, in terms of how it arouses other people’s interest and stimulates them, ‘cause that’s the way my experience is when I encounter some work of science, and I encounter lots of them, and they open my eyes, and I think ‘Oh my gosh how beautiful. Wow!’

#### Experiments and Iterations

We will refer to the prompt with Text 1 as Experiment 1, Text 2 as Experiment 2, and so forth. If the “experiment” prompt and the text to be analyzed were submitted repeatedly to the LLM during the same session (i.e., the user did not log out and then log back), we can expect that a similar or even identical output would be returned. However, this narrowing of responses would not reflect a general population trend but would more likely be an artifact of the initial conditions set when the LLM algorithm was initiated (Urbani, 2023). One important characteristic of large language models is that the underlying algorithms are designed to mimic variations within natural language processing. This difference depends on the random number captured by the LLM to set the initial conditions. As a result, the same prompt entered by different users (or different restarts through logging into the same account) will prompt different initial conditions into the LLM. It would be expected to return different responses. To take advantage of this effect, for each iteration, each time before we entered the text, we logged out of the LLM and then logged back in. These responses would eventually trend toward a similar result representative of the general population.

In our case, the LLM used in this analysis used a pseudo-random number generator (PRNG). It is this variation in output that takes advantage of the power of an LLM. While differences in interpretations between people are expected, an LLM uses the vast internet archive dataset harvested by the Common Crawl. This dataset offers a general interpretation of a population of many thousands and possibly hundreds of thousands of individuals when the same prompt is entered repeatedly. In our case, each time Experiment 1 text is entered by a different user, the responses may differ, but a trend in the responses was revealed over a series of these entries. We call each repeated entry of an experiment text an “iteration.” We performed 160 iterations for this study for each of the three experiments.

### Data Analysis Approach

For data analysis, we coded the responses from the LLM as either positive when the code was reported as present (i.e., coded as 1) or negative when the code was reported as absent (i.e., coded as 0). Sometimes, the response would be ambiguous about the presence or absence of a characteristic, so we applied a conservative interpretation of the data. Only entirely affirmative responses were coded as positive. If the response implied a level of uncertainty (e.g., when the LLM responded, “Some level of the characteristic is present.”), it was marked as negative.

#### Identifying an Outcome Measure

We recorded the results for each code in each experiment iteration. After 5, 10, 20, 40, 80, and 160 iterations, we calculated the proportion of positive responses for the given number of iterations. The results are shown in Tables 1, 2, and 3 in the Results and Discussion section. We will refer to this value as the large language model quotient (LLMq), where the total number of positive responses in a set of iterations was divided by the number of iterations for that given set. The equation is presented below:

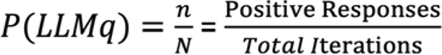

**Table 1:** Large language model quotients (LLMq) for five conceptual codes examining Text 1.

| Iterations | Autonomy | Persistence | Perception of identity | Novelty | STEM interests |
| --- | --- | --- | --- | --- | --- |
| 5 | 0.200 | 0.600 | 0.000 | 0.000 | 0.000 |
| 10 | 0.300 | 0.600 | 0.000 | 0.000 | 0.100 |
| 20 | 0.350 | 0.550 | 0.050 | 0.000 | 0.100 |
| 40 | 0.325 | 0.625 | 0.075 | 0.000 | 0.100 |
| 60 | 0.328 | 0.534 | 0.086 | 0.000 | 0.103 |
| 80 | 0.275 | 0.575 | 0.075 | 0.013 | 0.138 |
| 100 | 0.300 | 0.600 | 0.070 | 0.010 | 0.120 |
| 120 | 0.308 | 0.625 | 0.075 | 0.008 | 0.133 |
| 140 | 0.316 | 0.632 | 0.075 | 0.015 | 0.128 |
| 160 | 0.325 | 0.594 | 0.088 | 0.013 | 0.131 |

**Table 2:** Large language model quotients (LLMq) for five conceptual codes examining Text 2.

| Iterations | Autonomy | Persistence | Perception of identity | Novelty | STEM interests |
| --- | --- | --- | --- | --- | --- |
| 5 | 0.600 | 0.800 | 0.800 | 0.800 | 1.000 |
| 10 | 0.500 | 0.700 | 0.900 | 0.900 | 1.000 |
| 20 | 0.700 | 0.800 | 0.950 | 0.950 | 1.000 |
| 40 | 0.780 | 0.805 | 0.927 | 0.976 | 1.000 |
| 60 | 0.733 | 0.817 | 0.900 | 0.933 | 1.000 |
| 80 | 0.759 | 0.831 | 0.904 | 0.928 | 1.000 |
| 100 | 0.769 | 0.846 | 0.904 | 0.913 | 1.000 |
| 120 | 0.784 | 0.856 | 0.912 | 0.920 | 1.000 |
| 140 | 0.799 | 0.842 | 0.921 | 0.928 | 1.000 |
| 160 | 0.792 | 0.836 | 0.912 | 0.931 | 1.000 |

#### Comparison with Traditional Coding

The three sample interview excerpts were uploaded into a qualitative data analysis software package to facilitate data management and analysis.^1^ Three research team members carried out the traditional coding simultaneously and separately. None had access to the LLM coding results. All three coders are science educators and were not involved in collecting the interviews from which the quotes were extracted. These researchers only read and evaluated the selected quotes for the presence of the five conceptual codes used in this analysis. The coders used the same five codes included in the LLM prompts (i.e., Autonomy, Persistence, Perception of Own Identity or Self, Novelty, and *STEM Interests*). They assessed the transcripts line-by-line to identify text selection in the interview excerpts for the five codes. The coding results of the three traditional coders were compared, and any discrepancies or disagreements were discussed and resolved through group meetings.

## Results

### Calculating the Large Language Model Quotient

The LLM data analysis began by performing five iterations and then calculating the large language model quotient (LLMq) for this set of five iterations. Next, five iterations were performed, and the LLMq was calculated for 10 iterations. This sequence was repeated to produce LLMq values for 20, 40, 60, 80, 100, 120, 140, and 160 total iterations. These results for each experiment are shown in Tables 1, 2, and 3. The results were also graphed to display how the LLMq-values changed with each set of 20 total iterations. The results displayed in the graphs indicated that LLMq values varied for each of the three experiments across the five codes. However, it is essential to note that the LLMq values for a specific code within a specific experiment appeared stable with the trace of the graph leveling out.

**Table 3:** Large language model quotients (LLMq) for five conceptual codes examining Text 3.

| Iterations | Autonomy | Persistence | Perception of identity | Novelty | STEM interests |
| --- | --- | --- | --- | --- | --- |
| 5 | 1.000 | 1.000 | 1.000 | 1.000 | 0.200 |
| 10 | 1.000 | 1.000 | 1.000 | 1.000 | 0.200 |
| 20 | 1.000 | 1.000 | 1.000 | 1.000 | 0.350 |
| 40 | 1.000 | 1.000 | 1.000 | 1.000 | 0.317 |
| 60 | 1.000 | 1.000 | 0.983 | 1.000 | 0.333 |
| 80 | 1.000 | 1.000 | 0.964 | 1.000 | 0.325 |
| 100 | 1.000 | 1.000 | 0.971 | 1.000 | 0.317 |
| 120 | 1.000 | 1.000 | 0.976 | 1.000 | 0.312 |
| 140 | 1.000 | 1.000 | 0.978 | 1.000 | 0.302 |
| 160 | 1.000 | 1.000 | 0.981 | 1.000 | 0.327 |

### LLMq Results of Texts

For clarity, we offer the following examples of how LLMq values were calculated. For Text 1, concerning the concept of *Autonomy*, the first five iterations produced one positive response (See Table 1). In this instance, the LLMq was equal to 0.2, indicating that 20% of the prompted responses from the LLM were positive (i.e., the LLM stated the presence of the relevant code in the text). Next, for the set of 10 iterations with three positive responses, an LLMq value of 0.3 was calculated. For the third set, which consisted of 20 iterations, an LLMq value of 0.35 was calculated. These results and the subsequent calculations of LLMq values for the other four codes, *Persistence*, *Perception of Own Identity and Self*, *Novelty*, and *STEM Interests* are also displayed in Table 1. The results from Table 1 are graphed in Figure 1.

**Figure 1:**
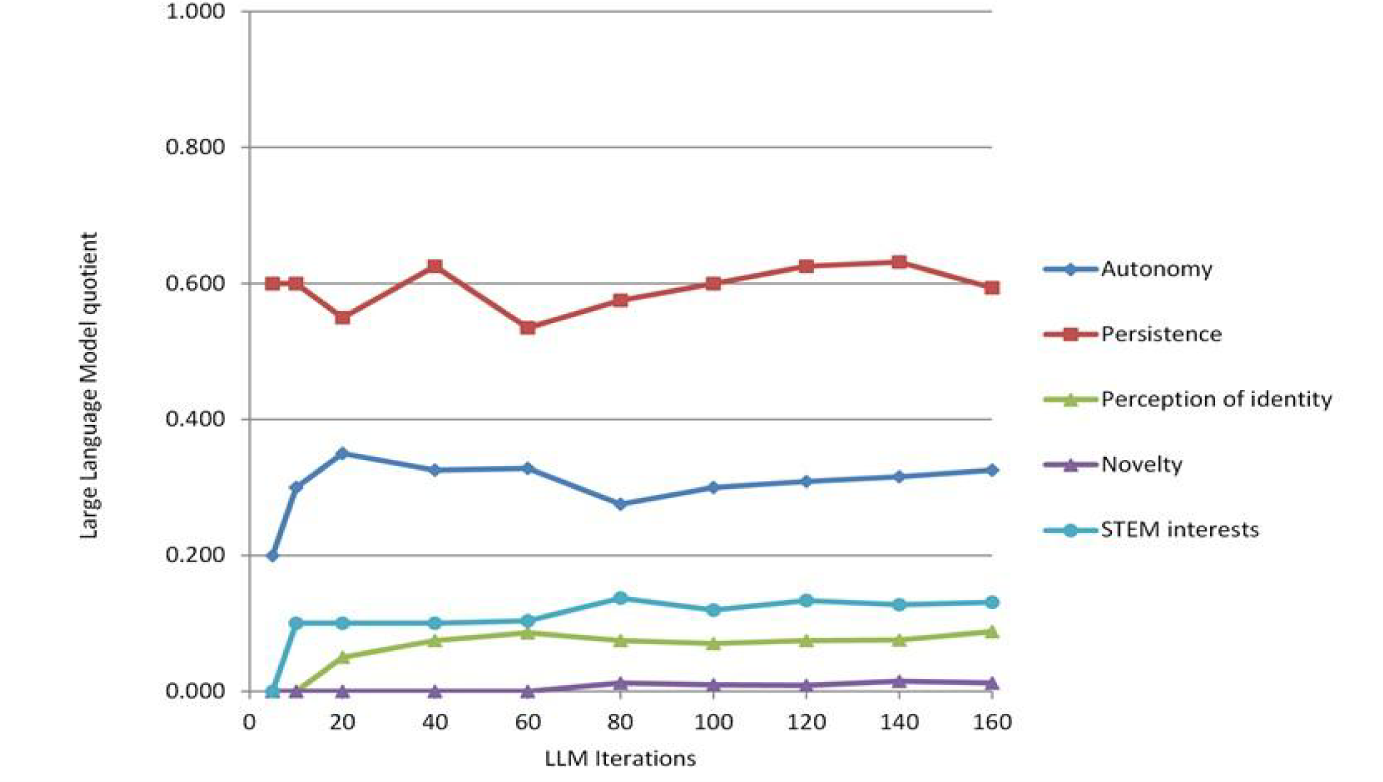
Graph of LLMq values shown in table 1 above

For Text 1, the LLMq results show low or very low levels of agreement across the five codes. Text 1 consisted of 76 words and was selected for this analysis for its brevity and general lack of precise detail with respect to the five codes considered in the analysis. For three codes, *Perception of Identity and Self*, *Novelty*, and *STEM Interests*, the LLMq results showed that they were identified in less than 15% of the iterations. For example, for *Perception of Own Identity and Self,* the LLM analysis results for 5 and 10 iterations, the LLMq was 0.000 or none of the iterations identified this code as present. As more iterations were carried out, the successive LLMq values were less than 0.100, which can be interpreted as the LLM identified this code in less than 10% of the iterations. More specifically, for 160 iterations, the result of LLMq = 0.088 was interpreted as the code *Perception of Own Identity or Self, which* is identified as present in only 8.8% of the iterations. The results for *Novelty* and *STEM Interests* may be similarly interpreted. The results for *Autonomy* showed that this code was identified as present by the LLM in up to 35% of the iterations. After 160 iterations, the results showed an LLMq = 0.325, indicating that *Autonomy* was identified as present by the LLM in 32.5% of the iterations. The results for *Persistence* showed that the LLMq values ranged from 5 iterations to 160 iterations, and the LLM identified this code in more than half of the iterations. At 160 iterations, the LLMq = 0.594 which indicates that the LLM reported this characteristic present in 59.4% of the iterations. Overall, the LLM analysis of Text 1 produced results indicating that none of the five codes were clearly present. The strongest result was for the code *Persistence* found by the LLM in about 60% of each set of iterations.

Next, we consider the results for the Text 2 LLM analysis. Here, the LLMq values for the codes *Perception of Own Identity or Self*, *Novelty*, and *STEM Interests* are 0.900 or greater across nearly all sets of iterations. The results also show that for the code *Persistence*, LLMq values were 0.800 or greater for all except one set of iterations. For the code *Autonomy*, LLMq values were 0.700 or greater for all except the initial two sets of 5 and 10 iterations.

Overall, the LLM analysis identifies all five of the codes to be present in most of the iterations. When the LLM identifies a code as present, the output included a response that in some cases cited selections from the text. Examples are shown in Figures 2A and 2B. Note that the LLM outputs are consistent, but are not identical.

**Figure 2:**
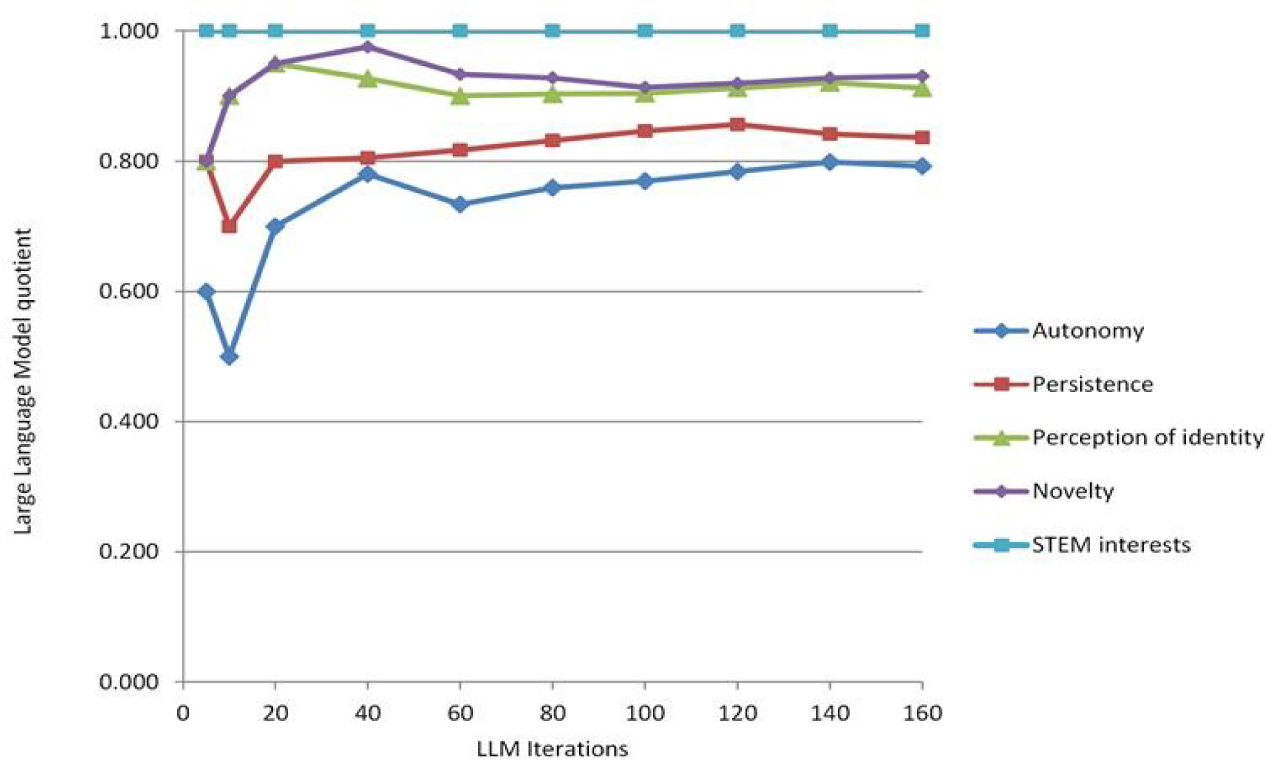
Graph of LLMq values shown in table 2 above

**Figure 2A:**
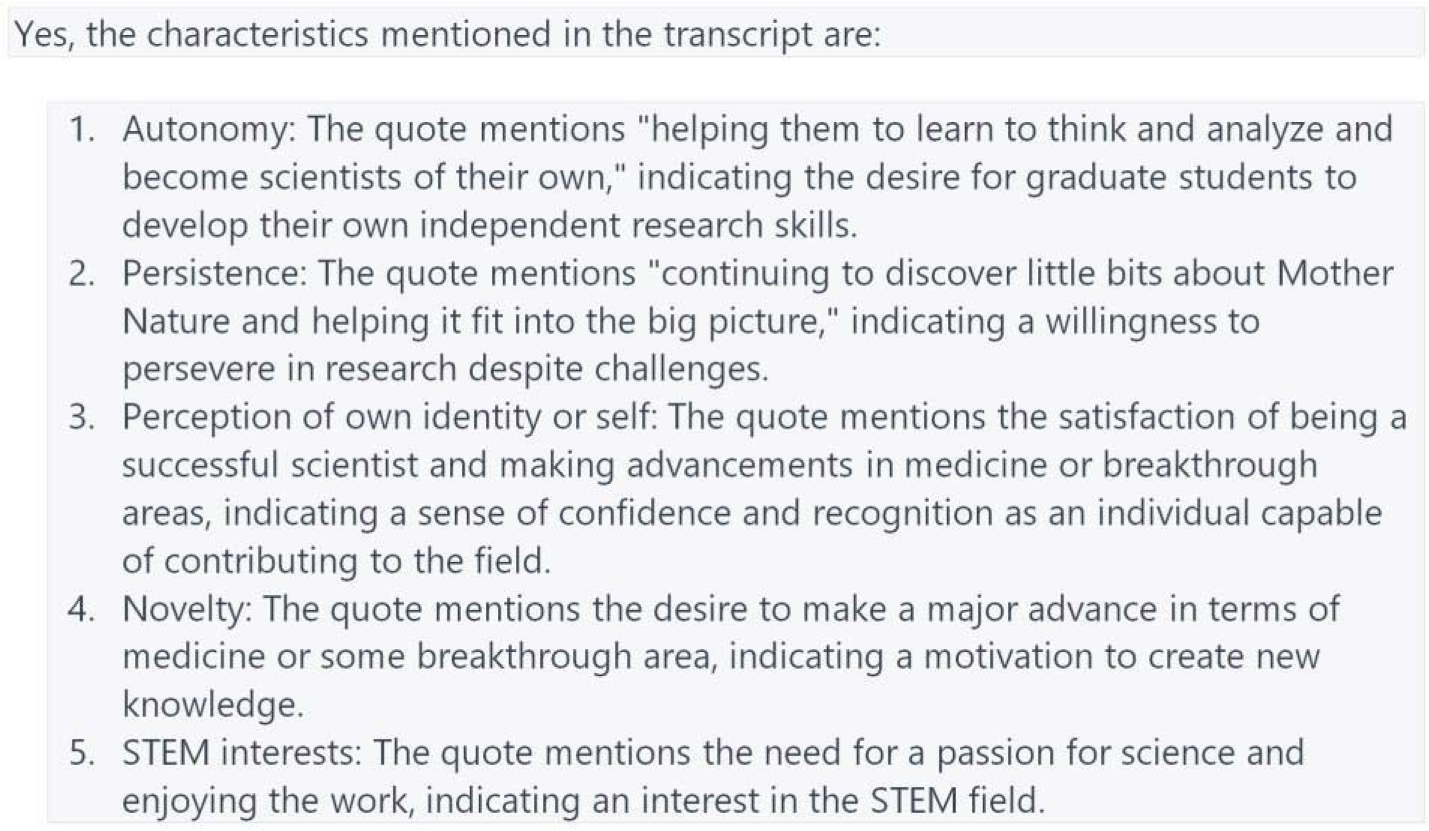
Example of LLM output for Text 2

**Figure 2B:**
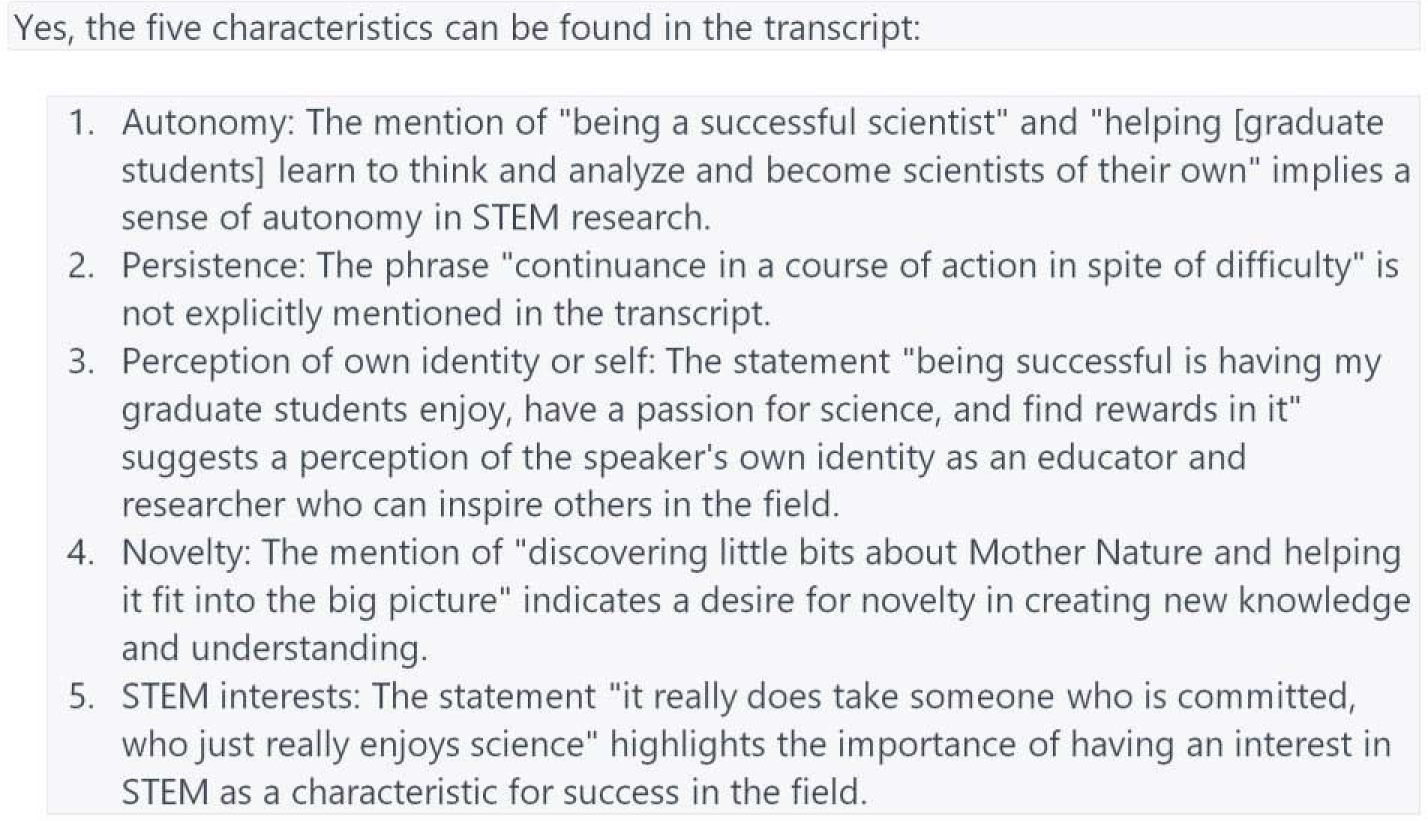
Example of LLM output for Text 2

Finally, the LLM analysis found that Text 3 had LLMq values that were definitive or nearly definitive for four codes: *Autonomy, Persistence, Perception of own identity or self, and Novelty*. However, for the code *STEM Interest*, the results were mixed. The LLMq values were stable as the graphed curve flattened out as iterations increased. The reported code *STEM Interest* was present in only one-third of the cases for 60 iterations and only 32.7% of cases for 160 iterations. (See Figure 3.) This result indicates that *STEM Interest* was not clearly present in Text 3.

**Figure 3:**
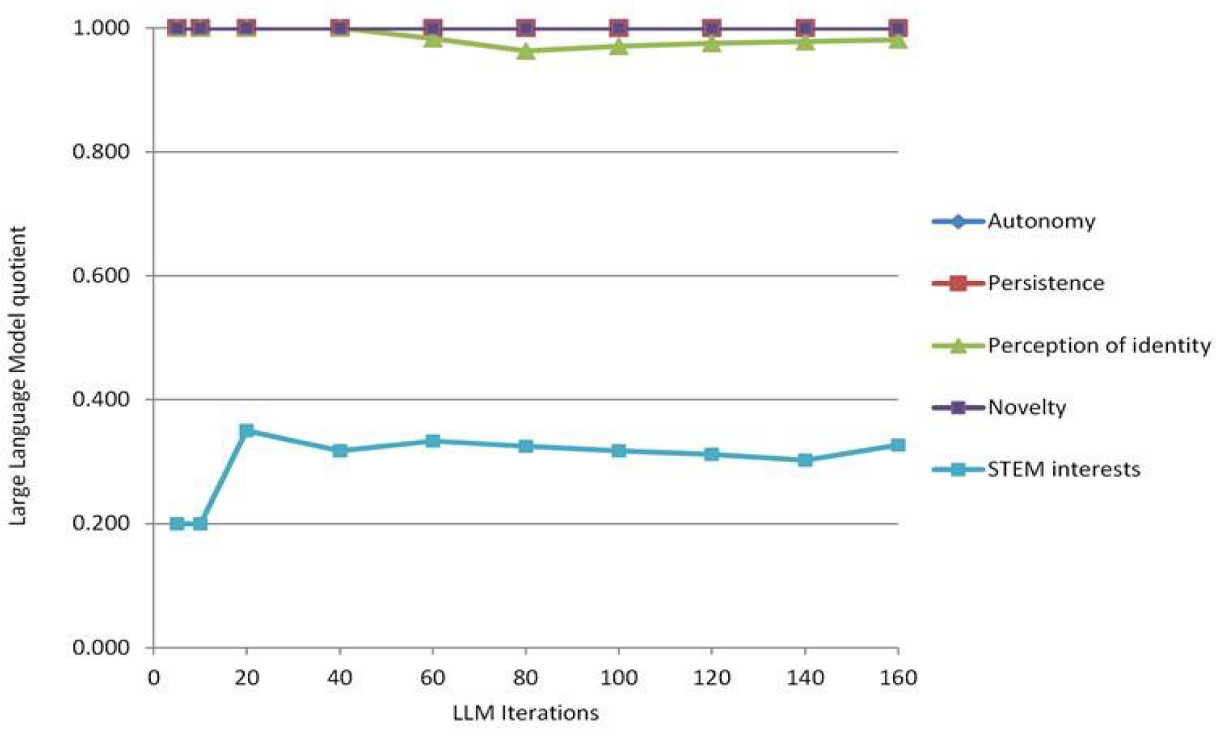
Graph of LLMq values shown in table 3 above

Two examples of LLM output responses for Text 3 are shown in Figures 3A and 3B. Comparing these two examples, the responses for the three codes, *Autonomy, Perception of Own Identity or Self,* and *Novelty*, are the same in the results of these two iterations of Experiment 3. However, the output identifies the presence of the *Persistence* code but cites two different selections from Text 3. Also, while both iterations do not identify the code *STEM Interest* in Text 3, the output in Figure 3B offers the following additional insight, “However, the overall context of the discussion revolves around STEM research, experiments, and encountering scientific work, indicating a focus on STEM interests.”

**Figure 3A:**
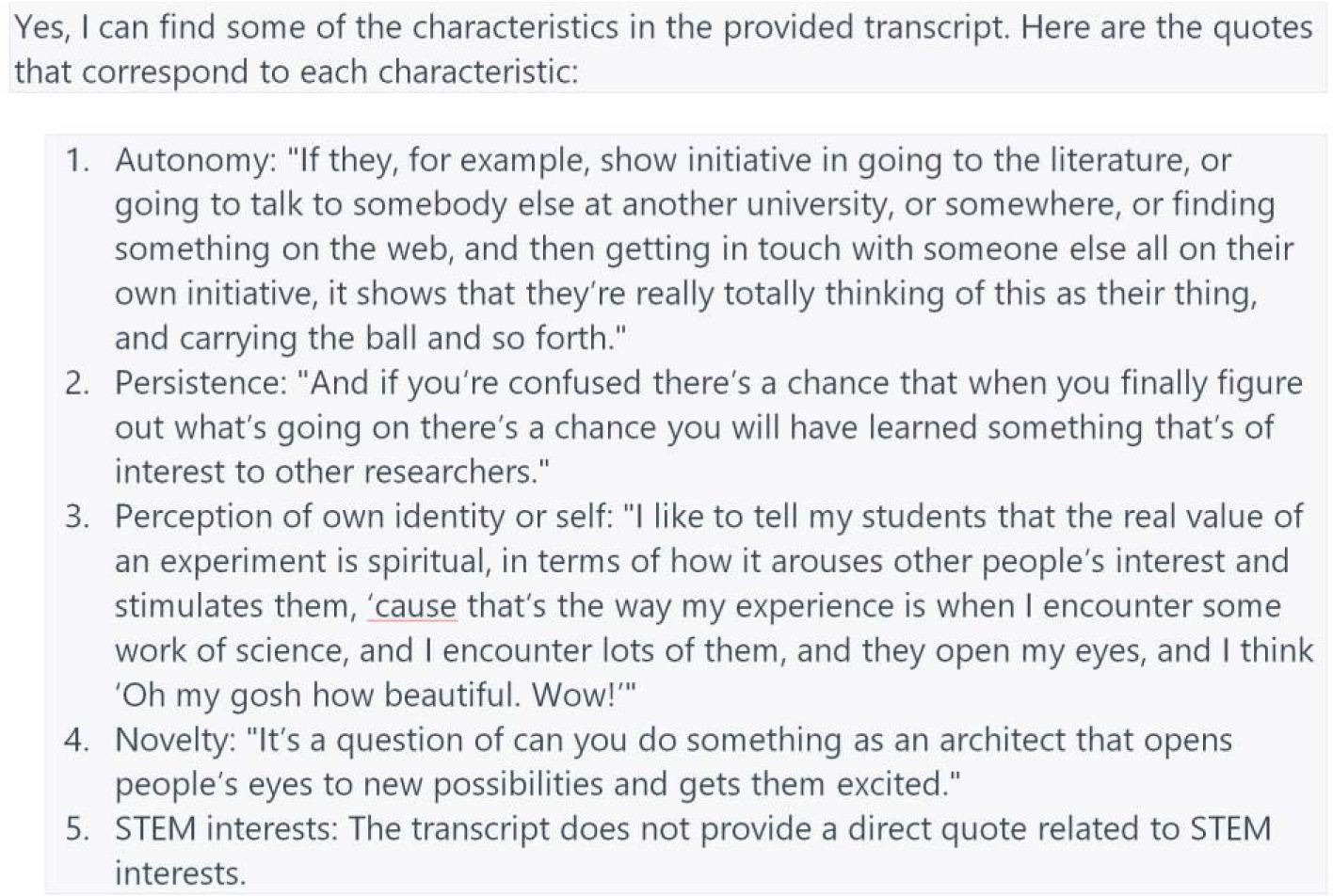
Example of LLM output for Text 3

**Figure 3B:**
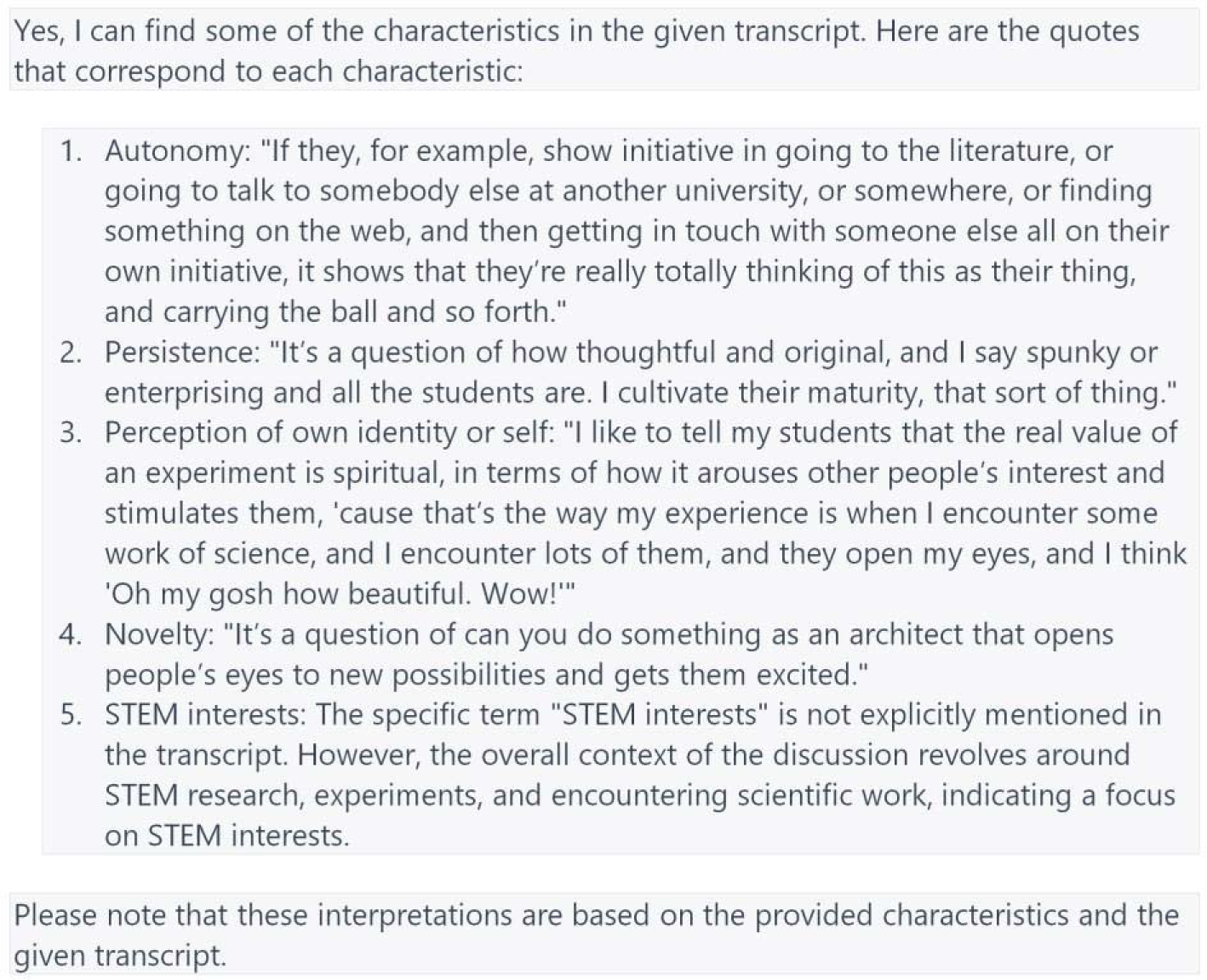
Example of LLM output for Text 3

### LLMq and Traditional Coding

As with the LLM analysis, traditional coding produced mixed results across the three narrative texts. In Text 1, the three human coders focused on the following Text 1 excerpt:

> *“ tried to be as independent as I could be and I did try a number of things, but then I would brainstorm with the postdoc and we met often with the advisor”*

Here, two coders cited this excerpt as an example of *Perception of Own Identity and Self*, while the third coder cited it as an example of *Autonomy*. Overall, the three coders did not find evidence for the codes *Persistence*, *Novelty*, or *STEM Interest* in Text 1. The results from the Text 1 analysis suggest that the five codes were either not present or were not clearly present. Comparing this result to the LLMq results, we find that the outcomes are similar across all five codes. The LLM textual analysis showed that none of the codes produced consistently positive results, the strongest results were for the code *Persistence* with an LLMq of about 0.600 or roughly 60 percent.

For Text 2, the results from the traditional coding found that all five codes were present. For example, the following excerpt was coded by two researchers as an example of *STEM Interests* and one researcher coded for *Perception of Own Identity or Self*.

> *“being successful is having my graduate students enjoy, have a passion for science”*

The following excerpt was coded by all three coders as demonstrating autonomy.

> *“analyze and become scientists of their own”*

An example of novelty was coded by all three coders below.

> *“ I think it would be wonderful to be able to really make a major advance in terms of medicine or some really breakthrough area, but for me it’s probably just discovering little bits about Mother Nature and helping it fit into the big picture”*

Next, the following excerpt was coded by all three researchers as demonstrating persistence.

> *“it really does take someone who is committed, who just really enjoys science”*

These results for Text 2 are consistent with the LLMq output for Text 2, where all five codes were present at above. 80 for the LLMq result.

For Text 3, traditional coding found examples of all five codes present in the text. This result is also consistent with the results from ChatGPT. For example, all three coders coded the following excerpt for *Autonomy*.

> *“If they, for example, show initiative in going to the literature, or going to talk to somebody else at another university, or somewhere, or finding something on the web, and then getting in touch with someone else all on their own initiative, it shows that they’re really totally thinking of this as their thing, and carrying the ball and so forth.*”

A second excerpt from Text 3 shown below was coded by two coders as *Novelty* and as both *Persistence* and *STEM Interests* by a third coder.

> *“And if you’re confused there’s a chance that when you finally figure out what’s going on there’s a chance you will have learned something that’s of interest to other researchers.”*

One coder identified the code *Perception of Own Identity or Self* in the following text, which matches the two LLM outputs cited above.

> *“I like to tell my students that the real value of an experiment is spiritual, in terms of how it arouses other people’s interest and stimulates them, ‘cause that’s the way my experience is when I encounter some work of science, and I encounter lots of them, and they open my eyes, and I think ‘Oh my gosh, how beautiful. Wow!’”*

When we compare the LLMq results to the results from traditional coding, there are areas of misalignment. These areas of divergence, specifically with the codes of *Perceptions of Identity or Self* and *Persistence* allowed the research team to have nuanced conversations about how we applied the codes in the analysis. For *Perceptions of Identity or Self*, human coders applied this code to texts highlighting self-descriptions of identity from the interviewed scientists. ChatGPT took the prompt it was provided and extended *Perceptions of Identity or Self* to include how scientists perceived themselves through other people. ChatGPT expanded the definition of identity to include how scientists view their self-identity as a result of the success of their students. This is a critical component to add to this particular code, and the research team would not have reflected on this without the external review of ChatGPT.

When examining *Persistence,* especially for Text 3, one human coder coded persistence, while LLMq showed that 100% of the iterations contained this code. ChatGPT identified the word “finally” as evidence of persistence from the text in its outputs. The research team re-evaluated the text. Based on the discussion, the research team realized that we were not carefully highlighting the word “finally” in the narrative. When we reflected on our code application, the research team agreed that Text 3 contained *Persistence* and re-coded the text to reflect this iteration of the code application.

The LLMq reported consistent and reliable results, which qualitative researchers can use to validate traditional coding. These results do not negate the importance of traditional coders but highlight the usefulness of the LLMq as a validation tool to identify and avoid potential implicit bias from researchers (Nosek et al., 2007). Similar to the practice of negotiation across coders when disagreements are identified, the LLMq results prompted the research team to expand the operative definitions of the codes.

## Discussion

This study explored using an LLM to perform qualitative coding analysis, leading to calculating the large language model quotient representing the percentage of positive outcomes returned from a set of analytical iterations. The results generated from the analysis of the iterations were compared to traditional coding and were found to be consistent. Recent work by Xiao et al. (2023) demonstrates the feasibility of using an LLM as “another rater” for qualitative analysis. However, in that study, only single LLM outputs were examined. Since LLMs are stochastic and are designed to generate variable responses from one output to the next, concluding a single LLM response is not the best approach. Multiple iterations of a prompt produce a better representation of LLM outputs. For this reason, we have chosen to explore the results across a range of output iterations for three different text examples.

Additionally, our results show that applying an LLM in performing qualitative data analysis produces consistent results with a sufficient number of repetitions. Gilardi et al. (2023) found similar results in which ChatGPT^®^ 3.5 produced accurate and reliable repeated classifications of Twitter posts regardless of randomness (temperature range 0.2-1). In their study, ChatGPT outputs were more reliable than traditional coders (>84%), and our results also confirm this finding.

An LLM can be used as a research instrument because it is designed to pull vast amounts of data and converge to a solution. This characteristic of LLMs works well with the considerations of qualitative research, which aims to identify meaning and trends within non-ordinal data. The LLMq values calculated for each of the three Texts examined in this analysis and graphed in Figures 1, 2, and 3 indicate that LLMq values appear fairly stable beyond 40 iterations. For each of the 5 codes, the trajectories of the graphed curves plateaued at this mark. In addition, the results from the traditional analysis are reflected in the LLMq values, even in the case of Text 1 for the code *Persistence*. Here, the traditional analysis showed a relatively low interrater agreement among the coders, and this lack of agreement is reflected in the LLMq values reported in Table 1 and Figure 1. The LLMq value after 160 iterations was 0.594 or 59.4% agreement from the LLM analysis that the code *Persistence* was found in Text 1. This inconsistent result from the LLM analysis was reflected in inconsistent traditional coding results. In this case, the LLMq values show that the LLM reports finding the code *Persistence* with a probability only 9.4% greater than random chance (50%). These results show that the LLM analysis was stable despite inconsistent results. These results offer some strong support for applying large language models as an analytical tool for qualitative data.

### Limitations

While LLM applications have shown great potential in aiding qualitative data research, they also have inherent limitations. The most significant limitation of the use of LLM is the algorithm itself. LLMs rely on patterns and structures present in the training data, and if specific linguistic nuances or subtleties are absent, the model’s understanding may be limited. The use of the data generated by the Common Crawl discussed earlier in this paper mitigates this limitation, drawing from the vast and growing body of content available through the internet.

The quality of the input data also plays a vital role in the effectiveness of LLMs in qualitative research. If the input data is of poor quality or contains biases or inaccuracies, the LLM’s outputs will likely reflect those shortcomings. Noisy data can also pose a challenge for LLMs in qualitative research. Noisy data refers to data that is unstructured, inconsistent, or contains anomalies such as transcription errors. LLMs may struggle to accurately interpret and assign codes to such data, leading to potential inaccuracies or misinterpretations. It is in these circumstances that hallucinations might occur. A careful review of transcripts can avoid this and is a common step in qualitative analysis.

The LLM limits the length of the input data. For example, at the time of data collection for this study, ChatGPT^®^ 3.5 had a default limit of 2048 characters or “tokens,” with an option to extend the length to 4096 tokens for a subscription fee. Each character, including spaces between words, is referred to as a token and is a measurement for the length of an inputted text. A longer text length allows for more extended quotes to be entered for each analysis. In certain instances, it may be necessary to enter quotes of an extended length to preserve contextual cues within the narrative that may figure into the LLM response. A quick back-of-the-envelope calculation indicates that 4096 characters amount to a few lines more than one page of single-spaced text.^2^

## Conclusions

These results offer some support that LLMs may be used as a tool to streamline qualitative research data analysis. Far from suggesting the replacement of researchers, it is clear that decisions about employing the LLM and interpreting its output are essential. LLMs offer researchers the autonomy to design prompts aligned with their interests and then test their ideas with an artificial intelligence engine designed to bring to bear massive data harvested from the internet. Creating a codebook-driven prompt may mimic traditional coding. In contrast, prompts that ask the LLM to generate a list of themes may lead to generating themes not initially considered by the researcher. Currently, LLMs are far from autonomous and should be prompted carefully to generate valuable data.

In this paper, we examined how the output of an LLM behaved when queried recursively across three different sets of texts using the same prompt. The results led to a simple representation of these results in the form of the quotient (LLMq). This simple calculation offers results that may be used for some helpful applications.

One potential application is using an iterative LLM analysis as a screening tool. Since the LLMq quantifies the degree of identification of a code within a text, entering a lengthy interview as a series of short excerpts would likely produce results highlighting sections of text where particular codes are present. Conversely, excerpts with low LLMq suggest a paucity of codes. While entering the type of prompt described in this paper would produce a binary outcome—that codes are either entirely present or absent in an excerpt— iteratively applying the prompt produces results with more nuance, asserting that codes may exist across a spectrum. Effectively, the LLMq optimizes the efficiency of researchers, who can hone in on areas of particular interest.

Another use of LLMq is as a post-hoc analysis tool. LLMq calculations can provide confirmatory or additional data to the traditional coding. The quantification of traits can justify the inter-rater reliability found, and perhaps the LLM may discover other traits not identified initially. A significant advantage of the LLMq is that a rater can independently and efficiently perform a “check” on their coding without relying upon other raters of equal competence. The LLM can provide supplementary or even new data to complement a researcher’s findings.

The future of LLM development holds immense promise for significant improvements in LLM performance that would increase the validity and reliability of this tool for qualitative research. As data sets used to train LLMs expand, there is an increasing focus on domain-specific and refined data. This data type will expose LLMs to a wider range of linguistic patterns and contextual information. This form of enhanced artificial intelligence training will enable LLMs to better reflect the nuances and complexities of qualitative data, leading to improved accuracy in code identification and interpretation. With access to more extensive and more diverse training data, future LLMs will be increasingly accurate in linguistic interpretations, thereby reducing the need for extensive iterations. This advancement will expedite the qualitative analysis process and facilitate more reliable and insightful findings. Researchers may use LLMs in an inductive process to search for themes and codes from the textual data. The application of an LLM as an inductive tool is beyond the scope of this current paper but is a topic for future investigation. By leveraging the power of future LLMs, researchers can unlock new possibilities for deeper understanding and exploration of qualitative data efficiently and accurately.

We envision the primary application of the LLMq would be to provide confirmatory data for a coding analysis. This approach offers individual researchers the option of using an LLM to gather confirmatory data to support their findings. Some readers might wonder at what level an LLMq might produce confirmation of a finding. To this question, we draw a parallel to a standard accepted quantitative value used to determine significance in a statistical finding, the probability value 0.05. A p-value of 0.05 indicates that the hypothesis under examination has a 5 percent chance of a finding being the result of chance. For our propose, an LLMq “cutoff” value of 0.950 or stated in other terms, a 95 percent positive detection rate of a particular code by the LLM. The 0.950 LLMq cutoff value is the equivalent of 19 out of 20 iterations returning an identification of the code by the LLM in the submitted narrative text. By leveraging LLM applications, researchers can augment their expertise while benefiting from multiple data sources to analyze transcripts and identify patterns and connections that might go unnoticed.

## Acknowledgement

Parts of this work have been supported by the US National Science Foundation (NSF DRL 1811265 and NSF REC 0440002). Any opinions, findings, conclusions, or recommendations expressed in this material are those of the author(s) and do not necessarily reflect the views of the US National Science Foundation.

## Research support

US National Science Foundation (NSF DRL 1811265 and NSF REC 0440002)

## Footnotes

1 The qualitative analysis software package used in this analysis was Dedoose^®^ (Los Angeles, CA).

2 Using the monospaced font, Monaco, at 12-point font, one single-spaced page has 54 lines at 65 characters per line for a one-page max of 3510 “tokens” per page.

